# Spatiotemporal control of microtubule acetylation by mechanical cues regulates lysosome dynamics at the immune synapse of B cells to promote antigen presentation

**DOI:** 10.1101/2023.08.28.555091

**Authors:** Felipe Del Valle Batalla, Isidora Riobó, Sara Hernández-Pérez, Pieta K Mattila, María Isabel Yuseff

**Affiliations:** Laboratory of Immune Cell Biology, Department of Cellular and Molecular Biology, Pontificia Universidad Católica de Chile, Santiago, Chile; Institute of Biomedicine, and MediCity Research Laboratories, University of Turku, Finland; Turku Bioscience, University of Turku and Åbo Akademi University, Turku, Finland; InFLAMES Research Flagship Center, University of Turku

## Abstract

The capacity of B cells to extract immobilized antigens through the formation of an immune synapse can be tuned by the physical characteristics of the surface where antigens are encountered. However, the underlying mechanisms that couple mechanosensing by B cells to antigen extraction and processing remain poorly understood.

We show that B cells activated by antigens associated with stiffer substrates exhibit enhanced spreading responses and higher tubulin acetylation at the center of the immune synapse, where less motile lysosomes preferentially localize. This process is coupled to the translocation of the microtubule acetylase, ATAT1 to the cytoplasm of B cells, which occurs as a mechano-response during BCR stimulation. Accordingly, B cells silenced for ATAT1 are unable to stabilize lysosomes at the synaptic interface and display a lower capacity to extract and present immobilized antigens to T cells.

Overall, these findings highlight how BCR-dependent mechano-responses trigger microtubule network modifications to precisely orchestrate lysosome positioning to promote antigen extraction and presentation in B cells.

## Introduction

Mechanosensing is a fundamental cellular process that enables cells to perceive and respond to physical environmental cues (Shaheen *et al*., 2019). This ability allows cells to adapt to tissue geometry and stiffness (also referred to as rigidity, measured in kPa), which in turn can influence genetic reprogramming, cell adhesion, migration, and organelle function (Chen *et al*., 2017). How mechanical cues associated with antigen recognition by B cells dictate their activation remains unresolved.

B lymphocytes interact with antigens that display diverse mechanical properties, ranging from less stiff immune complexes to highly rigid viral capsids (∼100 MPa) (Shaheen *et al*., 2019). Additionally, antigens can be recognized in a soluble form, considered to be in an environment with extremely low stiffness (0.01 kPa), or tethered to membranes of antigen-presenting cells (APC), which display stiffness values of approximately 0.1-0.5 kPa. B cells can also capture antigens associated with the extracellular matrix with rigidities of ∼20 kPa (Ciechomska *et al*., 2014). Moreover, the physical cues arising from membranes can change upon inflammatory conditions modifying the stiffness of the plasma membrane from the presenting cell (Bufi *et al*., 2015; Kim *et al*., 2019). Consequently, external physical cues emerge as an additional layer of information that B lymphocytes can sense and thereby impact their responses. Both B and T cells display membrane-bound immunoglobulin receptors, the BCR and T cell receptor (TCR), which possess the ability to sense the mechanical properties of their ligands (Zhu, Chen and Ju, 2019; Zhu *et al*., 2019). Additionally, B cells activated on stiff surfaces exhibit augmented BCR clustering and downstream signaling (Wan *et al*., 2013), leading to enhanced spreading responses and actin cytoskeletal rearrangements. Whether this has an impact on the capacity of B cells to extract and present antigens acquired at the immune synapse (IS) is unknown.

The mode of antigen extraction used by B cells depends on the physical properties of the substrate in which antigens are presented: antigens on flexible, softer surfaces are internalized into clathrin-coated pits by Myosin IIA-mediated pulling forces that trigger invagination of antigen-containing membranes (Hoogeboom and Tolar, 2016). During this process, the complexes formed by the B cell receptor and the antigen (BCR-Ag complexes) are internalized at regions enriched in actin foci (Roper *et al*., 2019). On the other hand, antigens presented on more rigid surfaces require the local fusion of lysosomes at the synaptic membrane that release proteases and acidify the synaptic cleft, facilitating antigen extraction (Yuseff *et al*., 2011). This proteolytic mode of antigen extraction relies on the precise positioning, tethering and secretion of lysosomes at the IS. Various factors regulate this process, including local actin cytoskeleton remodeling by proteasome activity (Ibañez-Vega, Del Valle Batalla, *et al*., 2019), proper assembly of the exocyst complex (Sáez *et al*., 2019), and the v-Snare VAMP7 to the plasma membrane (Obino *et al*., 2017).

Both cytotoxic T lymphocytes and NK cells enhance perforin-mediated killing at the immune synapse when interacting with antigens associated to stiff substrates (Basu *et al*., 2016), suggesting that lytic granule secretion is coupled to mechanosensing (Friedman *et al*., 2021). However, the underlying mechanisms behind this process remain largely unknown. In other cell types, the rigidity of the environment regulates membrane trafficking through integrin and RhoA-dependent remodeling the actin cytoskeleton to promote docking of secretory vesicles (Lachowski *et al*., 2022; Phuyal *et al*., 2022). Notably, the role of mechanosensing in directing lysosome positioning has not been previously characterized and B cells emerge as a valuable model to study this process.

Vesicles move along microtubules (MT) by their association to motor proteins, which can be regulated by post-translational modifications (PTM) of the microtubule network (Pu *et al*., 2016). In neurons, acetylation of microtubules fine-tunes the trafficking of lysosomes by modifying their affinity with kinesins (Morelli *et al*., 2018). In T cells, upregulation of tubulin acetylation by inhibition of the deacetylase HDAC6 leads to impaired transport of lytic granules to the synaptic membrane interacting with the target cell, highlighting how microtubule PTM, in particular acetylation, impact vesicle transport (Núñez-Andrade *et al*., 2016).

Thus, mechano-responses can regulate MT posttranslational modifications of the microtubule network, resulting in differential trafficking of vesicles to sites where membrane exchange or exocytosis is required. Interestingly, acetylation of tubulin can also be coupled to mechano-responses and is significantly enhanced at the leading edge of migrating fibroblasts when traction forces occur, promoting the release of GEF-H1 and acto-myosin contractibility (Seetharaman *et al*., 2022). In this work, we show that activation of B cells on stiffer substrates enhances the concentration of lysosomes at the center of the IS and ultimately improves their capacity to extract and present antigens. This process relies on increasing levels of tubulin acetylation in response to stiffness and the translocation of the acetylase ATAT1 from the nucleus to the cytoplasm. Elucidating how physical properties of antigen presenting surfaces are coupled to B cell effector responses that drive antigen extraction, processing and presentation, may provide valuable therapeutic tools for vaccine design, drug delivery or treatment of autoimmune diseases.

## Materials and methods

### Cell lines, culture, and treatments

In this study we used the mouse A20 lymphoma cell line, an FcγR-defective B cell line with the phenotype of quiescent mature B-cells, and the LMR7.5 Lack T-cell hybridoma, which recognizes I-Ad-LACK156–173 complexes. Both cell lines were cultured as previously described (Sáez *et al*., 2019) in CLICK medium (RPMI 1640, 10% fetal bovine serum, 100U/mL penicillin-streptomycin, 0.1% β-mercaptoethanol, and 2% sodium pyruvate).

For the induction of tubulin acetylation, cells were treated with 1 uM of the microtubule-stabilizing agent SAHA (TOCRIS bioscience Cat. No. 149647-78-9) for 30 minutes before activation or experimental procedures. For time-lapse acquisitions, lysosomes were labeled with Lysotracker DN-99 Red (Thermo Scientific, L7528).

### Preparation of tunable-stiffness poly-acrylamide (PAA) gels

For preparing the tunable-stiffness PAA gels, two sets of coverslips are used: a silanized-bottom coverslip where the PAA gel is immobilized and an smaller impermeabilized coverslip on top to create a “sandwich” to allow for the polymerization of the PAA gel (Charrier *et al*., 2020). Bottom coverslips are activated with a 2% 3-Aminopropyltrimethoxysilane (3-APTMS) (Sigma Aldrich 13822-56-5) in 96% ethanol solution for 5 minutes and washed with 70% ethanol. The top coverslips are siliconized or waterproofed with Rain-X. The coverslips are then washed with ultrapure water, dried well, and moved to the next stage.

The following reagents were used to prepare the PAA gels: 2% bis solution (Bio-Rad #1610142), 40% acrylamide (Bio-Rad #1610140), 10% APS in 10 mM HEPES solution, and TEMED (Table 1).

**Table 1:** preparation of poly-acrylamide gels of 0.3 (soft) and 13 (stiff) kPa.

| kPa | 40%<br>Acrylamide<br>(μL) | 2%Bis-Acrylamide<br>(μL) | 10% APS<br>(μL) | TEMED<br>(μL) | 1X PBS<br>(μL) | Total<br>(μL) |
| --- | --- | --- | --- | --- | --- | --- |
| 0.3 | 75 | 30 | 10 | 1 | 895 | 1011 |
| 13 | 233,75 | 125 | 6,25 | 1,8 | 883 | 1249 |

To prepare the gel, 9 uL of the gel solution was pipetted onto the center of the bottom coverslip. Then, it was well covered with the smaller cover and pressed gently until the mix spread outwards. After 30 minutes of polymerization, they were covered with aluminum foil. Next, 1X PBS was added to carefully remove the top cover. The gel can be stored in fresh 1X PBS at 4°C overnight and used within 48 hours.

For conjugation with ligands a solution of 0.5 mg/mL SULFO-SANPAH (Pierce, Thermo-Scientific #A35395) in HEPES buffer 10 mM was prepared. PBS was removed from the gels and immediately coated with SULFO-SANPAH at room temperature (RT). Gels were exposed to 365 nm UV light (Maestrogen UV illuminator MLB-16) for 10 minutes and washed with 1X PBS thrice. Finally, a BCR^+^ ligand (F(ab′)2 goat anti-mouse IgG) (Jackson ImmunoResearch, 115-006-146) was added to coat the gels at a concentration of 0.13 mg/mL and incubated overnight. This was followed by 3 washes with 1X PBS and immediately used for experiments or stored at 4°C protected from light for no more than 48 hr.

### Activation and immunofluorescence of B cells on PAA gels

80 μL of B cells (1.0 x 10^6^ cells/mL in CLICK medium with 5% FBS) were seeded onto an antigen (BCR ligand^+^)-coated gel for different time points in a cell incubator at 37 °C / 5% CO2. After each time point, the media was carefully aspirated off each PAA gel, and 100 μL of cold 1X PBS was added to stop the activation. PBS was removed, and each PAA gel was fixed with 50 μL of 3% PFA for 10 min at RT. PAA gels were washed three times with 1X PBS. The 1X PBS was removed, and 50 μL of blocking buffer (2% BSA and 0.3 M glycine in 1X PBS) was added to each coverslip.

Primary antibodies were diluted in permeabilization buffer (0.2% BSA and 0.05% saponin in 1X PBS) and incubated by adding 40 uL over the gels in a humid chamber at 4 °C overnight. The plate was sealed to avoid evaporation of the antigen solution. Gels were washed three times with permeabilization buffer. Secondary antibodies or dyes were diluted in permeabilization buffer, using 40 μL per PAA gel, and incubated for 1h at RT in dark and humid chambers. The PAA gels were washed twice with permeabilization buffer and once with 1X PBS. The PBS solution was removed from the coverslips. 8 μL of mounting reagent was added to a microscope slide. The PAA gels were mounted onto the slide with the cell side facing down. The slides were allowed to dry for 30 min at 37 °C or RT overnight protected from light.

### Immunoblot

Cells were lysed with 40 uL of RIPA buffer, then supernatants of samples were collected and loaded onto gels and transferred onto polyvinylidene fluoride membrane (Trans-Blot Semi-Dry Transfer Cell; Bio-Rad). Membranes were blocked in 2%BSA/TBS + 0.05% Tween-20 and incubated overnight at 4 °C with primary antibodies, followed by 60-min incubation with secondary antibodies. Western blots were developed with Westar Supernova substrate (Cyanagen, Cat. No. XLS3,0100), and chemiluminescence was detected using the iBright imager (Thermo Fischer).

### Atomic Force Microscopy (AFM)

The elastic modulus of the PAA gels (Young’s modulus, E) was assessed using a JPK NanoWizard with a CellHesion module mounted on a Carl Zeiss confocal microscope, Zeiss LSM510 (AFM; JPK instruments), and silicon nitride cantilevers (spring constant: 1 Nm^−1^, spherical 10 μm diameter tip; Novascan Technologies). The cantilever spring constant and deflection sensitivity were calibrated in fluid using the thermal noise method (Hutter and Bechhoefer, 1993).

Force measurements were then conducted at different locations (0.5 mm apart in x and y coordinates) within the region of interest. In each location, nine indentations distributed in a 3×3 point grid (30 μm × 30 μm) were performed. The elastic modulus for each force curve was calculated using JPK data processing software (JPK DP version 4.2), assuming a Hertz impact model.

### Antibodies & dyes

For primary antibodies, we used Rat anti-mouse LAMP1 (BD Bioscience, #553792), Rabbit anti-mouse Acetyl-Tubulin (Lys40, D20G3) (Cell Signaling, #5335), Rabbit anti-mouse αTubulin (Abcam, #ab6160), Goat anti-mouse IgM Fab2 (Jackson ImmunoResearch), Rabbit anti mouse GEF-H1 (Abcam #ab155785), Rabbit anti-mouse ATAT1 (Thermo Scientific, #PA5-114922), Rabbit anti-mouse YAP (D8H1X) XP (Cell Signaling #14074). For secondary antibodies: Donkey anti-rabbit IgG-Alexa 488/546/647 (Jackson Immunoresearch, 711-546-152), Donkey anti-rat IgG-Alexa 488/546/647 (Jackson Immunoresearch, 712-166-153), Phalloidin Rhodamine (Thermo Scientific, R415), Lysotracker-red DND 99 (Thermo Scientific, L7528) and Hoescht (Abcam, #ab228551).

### Cell transfection and electroporation

The Nucleofector R T16 (Lonza, Gaithersburg, MD) was used to electroporate 4 x 10^6^ A20 B cells with 3 μg of plasmid DNA using the LC-013 program. After transfection, cells were cultured for 18 ±2 hrs at 37 °C and 5% CO2 before functional analysis. For ATAT1 silencing, cells were electroporated with 100 nM of siRNA (sc-108799-SH, Santa-cruz technology) or 100 nM of siRNA control (sc-37007, Santa-cruz technology).

### Antigen presentation assay

Antigen presentation assays were performed as previously described on (Yuseff *et al*., 2011) with modifications. Briefly, B cells were incubated with either Lack-BCR-Ligand or BCR Ligand coated PAA gels and PAA gels with different concentrations of Lack peptide (Lack 156-173) for 1h. Then cells were washed with PBS and incubated with Lack-specific LMR 24 7.5 T Cells in a 1:1 ratio for 4h in a cell incubator at 37°C and 5% CO2. Supernatants were collected, and interleukin-2 cytokine production was measured using BD optiEA Mouse IL-2 ELISA set following the manufacturer’s instructions (BD Biosciences, Cat no. 555148).

### Cell imaging and image analysis

For widefield imaging, all Z-stack images were obtained with 0.3 μm between slices. Images were acquired in a widefield - epifluorescence microscope (Nikon Ti Eclipse) with an X60/1.25NA objective. For confocal acquisition, images were obtained in a Zeiss LSM880 Confocal with Airyscan detection microscope using a 63X/1.4NA oil immersion lens, with a Z-stack configuration of 0.2 μm. All acquisition for experiment replicates were performed with same mW/cm^2^ intensity for all illumination or equivalent exposition. The images were processed using Zeiss Black Zen software for Airyscan processing and analyzed with Fiji (ImageJ) (Schindelin *et al*., 2012).

For image analysis, when quantifying cell spreading a threshold mask (Otsu) was used in parallel with manual segmentation in instances when automatic algorithms were not able to resolve general actin structures. For actin foci quantification, we performed analysis of particles that were found inside previously detected spreading cells. In this case, we took into account workflows that have been tested for determining such structures (Roper *et al*., 2019). Other spot segmentation and cluster determination algorithms such as lysosome number at the IS, their size and pFAK counts were carried out with conservative algorithms of background subtraction and the same strategies of segmentation for each experimental set. Lysosome dynamics was studied with Fiji’s TrackMate plugin using the LAP tracker in settings recommended for its unsupervised setting, always testing for quality score of tracks. Overlap coefficients for colocalization and fluorescence ratios were determined by quantification of the mean fluorescence intensity of segmented structures and for single planes of acquisition at the IS plane (closest z plane to the PAA gel surface). Specifically, for colocalization analysis only single planes and individual regions of interest of the single cells were used and quantified with JaCoP plugin for Fiji software following its instructions. Determination of the localization index for differential accumulation of lysosomes in central or peripheral zones of the cell, was adapted from previous publications (Ibañez-Vega, Fuentes, *et al*., 2019). Nuclear and cytoplasmic accumulation was calculated following “Cyt/Nuc” plugin for Fiji (Grune *et al*., 2018).

### Statistical analysis

All data presented was tested for normality and homoscedasticity; accordingly, appropriate statistical tests were applied considering those factors. Data in plots and graphs are expressed as fold change of mean or mean ± SEM. If corresponding, experiments were analyzed by Student’s t-test or Mann–Whitney test following Gaussian and non-Gaussian distribution, respectively after performing d’Agostino and Pearson omnibus normality test. In cases where two conditions are shown, two-way ANOVA with Sidak’s multiple comparison test was performed if data followed normal distribution. When datasets followed non-gaussian distributions, Kruskal-Wallis tests were applied with Dunn’s a-posteriori multi comparison examination. Experiments were carried out with a sum of n ≥ 30 cells pooled from N = 3 biological replicates. Error bars shown are mean ± SEM. Statistical analysis was performed with Prism (GraphPad Software) and RStudio. For significance, p-values were calculated using different tests mentioned above and are illustrated in each figure.

## Results and discussion

### The rigidity of the antigen-associated surface regulates actin remodeling and spreading responses in B cells

Upon interaction with surface-tethered antigens, B cells trigger a spreading response to maximize antigen encounter and uptake (Yuseff *et al*., 2013). In this study, we aimed to characterize cytoskeletal and vesicular responses coupled to mechanical cues sensed through the B cell receptor (BCR).

To examine this, B cells were activated for different time points on surfaces containing BCR ligands associated to polyacrylamide (PAA) gels of 0.3 and 13 kPa, hereafter referred to as "soft" and "stiff" gels, respectively (Fig. EV1 A). The stiffness of the gels was measured by atomic force microscopy (AFM) and values obtained for soft and stiff gels (Fig. EV1 B) were consistent with previous reports. These mimicked physiological and pathological mechanical cues (Wan *et al*., 2013). BCR^+^ ligands (F(ab′)2 anti-IgG) coupled to both soft and stiff gels exhibited similar binding densities (Fig. EV1 C) and formed a continuous and homogeneous layer under all conditions.

B cells seeded onto gels of different stiffness, without BCR^+^ ligands exhibited no alterations in cell spreading after 5 and 30 minutes (Fig. EV1 D-E). Conversely, B cells seeded over stiff substrates containing BCR^+^ ligands, exhibited higher spreading responses after 15 and 30 minutes, in comparison with those seeded over softer substrates (Fig. 1A). Mean areas of B cells activated under stiff conditions were in the range of 300-400 μm^2^ compared to an average of 100 μm^2^ for B cells activated under soft conditions (Fig 1B). These observations suggest that B cells possess a BCR-dependent mechanosensing pathway that controls cell spreading in response to substrate rigidity coupled to BCR^+^ ligands.

**Fig 1.**
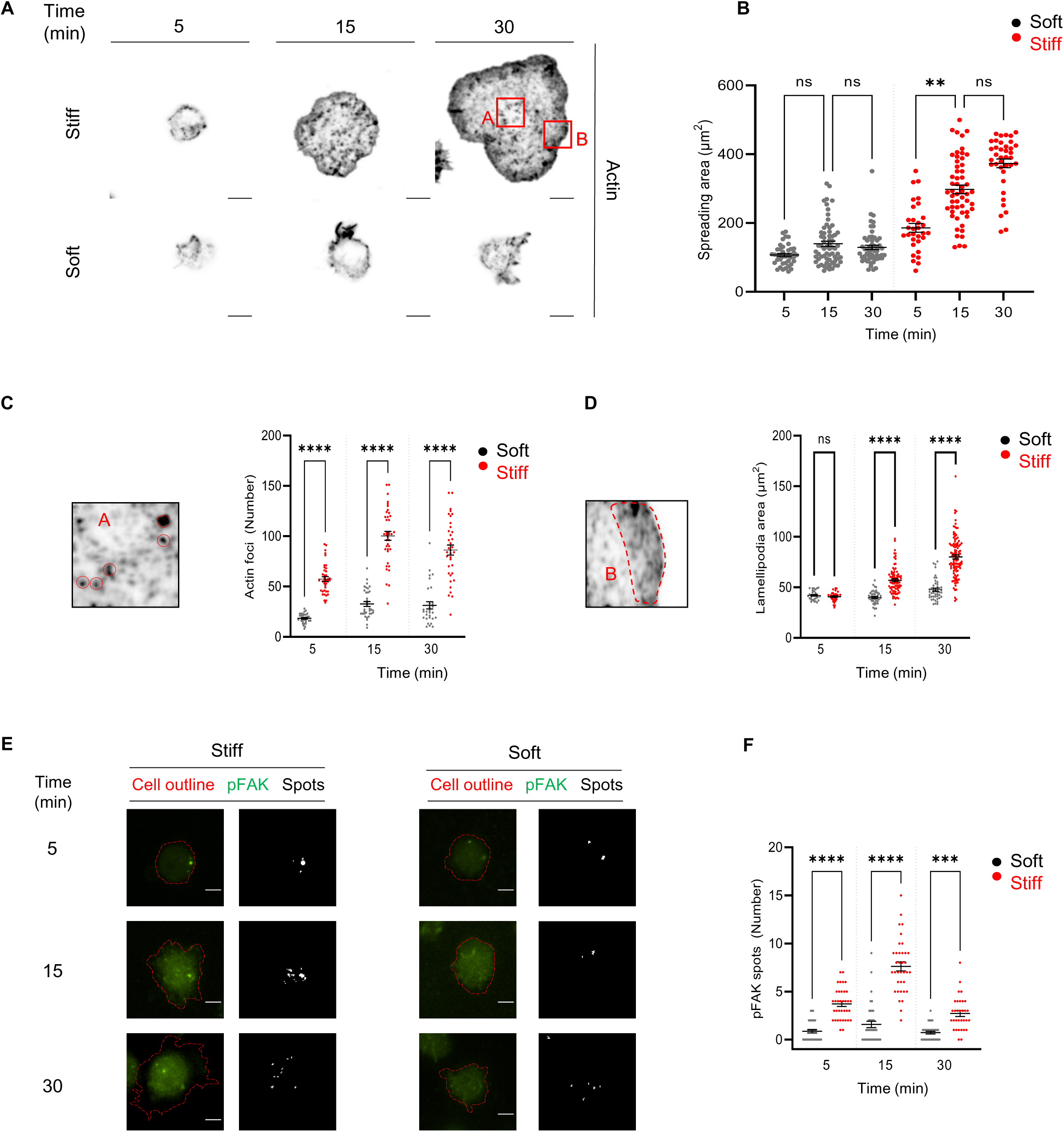
Substrate stiffness regulates cytoskeleton B cell responses upon BCR stimulation (A) B cells seeded over stiff or soft substrates at different activation times. Actin (phalloidin) is shown in black. Red squares delimit actin foci and lamelipodia areas. (B) Quantification of cell spreading areas in (A) defined by actin staining. (C-D) Quantification of actin foci detected and lamellipodia area, respectively, for cells seeded on soft or stiff substrates. (E) Representative images of fixed cells interacting with soft or stiff substrates. pFAK in green and cell outline in red. pFAK is shown as segmented particles (spots). (F) count of pFAK spots from experiments performed in (E). For every experiment, shown data considers n ≥ 30 cells pooled from N = 3 biological replicates. All scale bars are 5 μm. P values illustrated with asterisks are ** < 0.01,*** <0.001, **** <0.0001. Error bars are mean ± SEM.

Additionally, when examining actin cytoskeleton organization in these cells, we observed an increase in the number of actin foci and lamellipodia area (Figs. 1C, D insets A and B respectively) in B cells activated on stiffer substrates compared to soft substrates. Actin foci structures are considered a hallmark of B cell mechano-responses by engaging on inward forces that facilitate antigen concentration and internalization (Roper *et al*., 2019). These results support the idea that mechanical cues shape the activation of B cells, impacting the structure of the actin cytoskeleton, prompting us to explore how B cells couple the detection of physical cues during B cell activation to regulate antigen extraction and presentation.

### B cells elict mechanotransduction pathways during activation as a response to variable substrate stiffness

Until now, our results reveal that during activation, B cells remodel their actin cytoskeleton in response to external physical cues. The focal adhesion kinase (FAK) is involved in mechanosensitive responses, promoting the assembly of focal adhesions to support membrane protrusion during cell spreading and migration (Li *et al*., 2023). In B cells the phosphorylated active form of FAK (pFAK) is increased upon BCR stimulation in an integrin-independent manner known as “inside-out activation” through phosphorylation by PKCb (Shaheen *et al*., 2017). Thus, we next sought to determine whether in our model B cells could promote FAK activation in response to substrate stiffness. Our results show that B cells activated on stiff surfaces display increased levels of pFAK, observed as individual spots, in comparison to soft conditions (Fig. 1E-F). pFAK spot number was upregulated at earlier time points of activation (5 to 15 minutes) and decreased after 30 minutes. We confirmed this by immunoblot analysis, which showed higher levels of pFAK in cell lysates from B cells activated on stiffer substrates at similar time points (Fig. EV1 F-G).

We also verified that B cells induce stronger BCR signaling responses when activated on stiffer versus soft substrates (Fig. EV2 A). After 15 minutes of activation, B cells activated on stiffer substrates displayed higher levels of phospho-AKT and phospho-ERK in comparison to softer conditions (Fig. EV2 B). Taken together, these findings show that BCR-mediated mechanosensing in response to substrate stiffness activates FAK and BCR downstream signaling molecules that regulate spreading and signaling pathways associated with enhanced states of B cell activation.

### Tubulin acetylation is enhanced in a mechanosensory-dependent manner during B cell activation

During engagement with antigens, B cells remodel their actin cytoskeleton according to external mechanical cues; however, the impact of such cues on lysosome trafficking have not been addressed.

Intracellular trafficking relies on the organization of the MT network, where posttranslational modifications of tubulin can tune MT dynamics and impact vesicle transport (Janke and Magiera, 2020). Acetylation of alpha tubulin at lysine 40 stabilizes the tubulin lattice, and ensures the accumulation of KIF proteins that promote synaptic vesicle transport in neurons (Bhuwania, Castro-Castro and Linder, 2014). Whether physical cues regulate lysosome transport at the IS by modifying microtubule PTMs remained to be determined. Thus, we evaluated if acetylation of the MT network is regulated by physical cues sensed by B cells through their BCR. To this end, B cells were seeded over soft or stiff BCR^+^-coupled PAA gels for 5, 15 and 30 minutes and stained for alpha tubulin (a-Tub) and acetylated tubulin (Ac-Tub). Noticeably, B cells activated on stiff surfaces displayed higher levels of Ac-Tub compared to cells activated on soft substrates (Fig. 2A), which was measured as the total amount of Ac-tubulin and as a ratio to total tubulin levels (Fig. 2 B-C). This result was confirmed by western blot analysis of cell lysates from B cells activated on stiff and soft substrates, showing that levels of Ac-Tub were upregulated in a time and stiffness-dependent manner (Fig EV2 C-D).

**Fig. 2.**
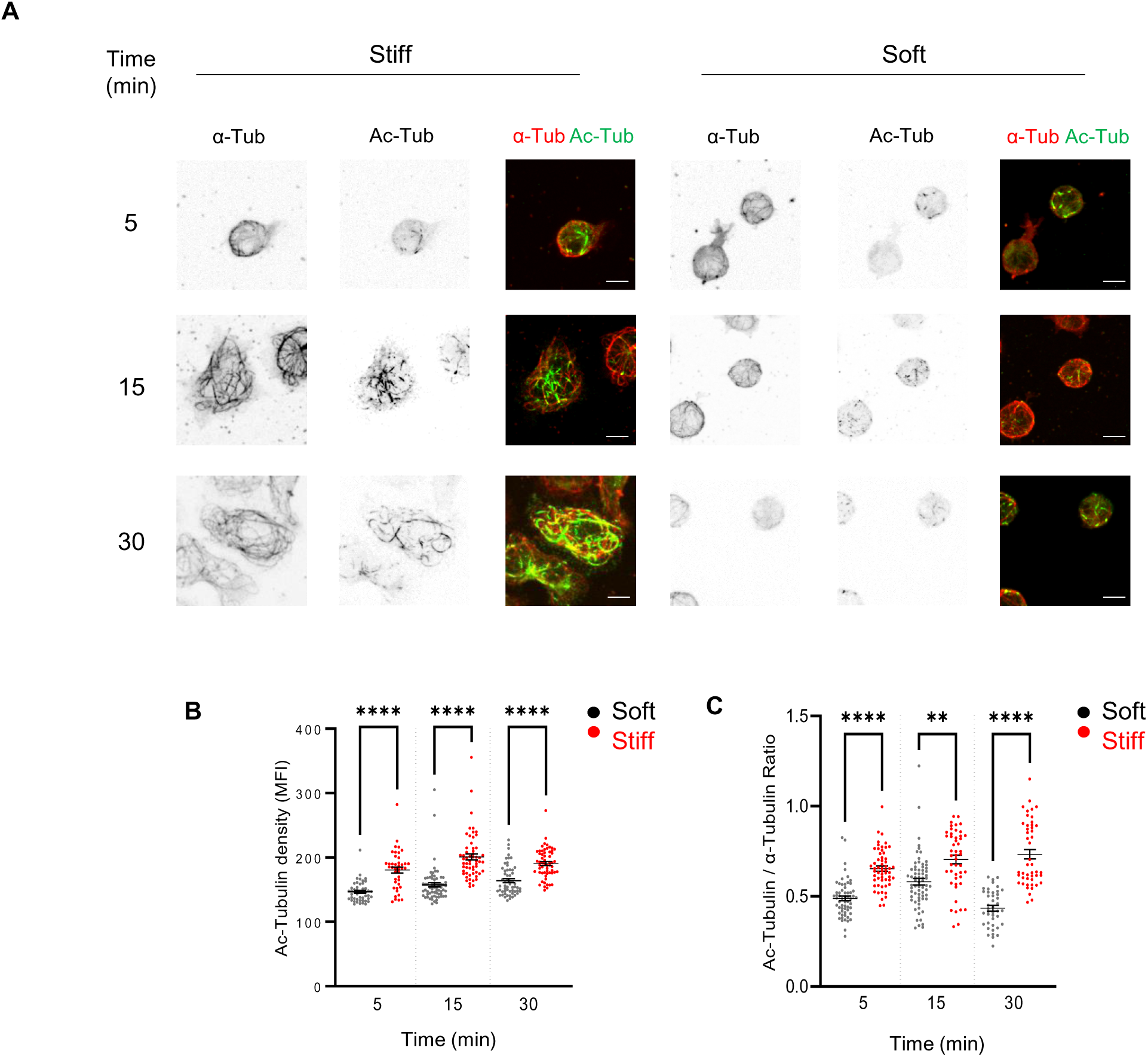
Tubulin acetylation is enhanced during B cell activation on stiff substrates (A) Representative images of fixed cells interacting with soft or stiff substrates at different time points. Alpha tubulin (a-Tub) and Acetylated tubulin (Ac-Tub) are shown separately and in composite format. (B) Ac-Tub density calculations based on MFI at the IS for images in (A). (B) Ac-Tubulin / α-Tubulin MFI ratios quantified at the IS for images in (A). For every experiment, shown data considers n ≥ 30 cells pooled from N = 3 biological replicates. All scale bars are 5 μm. P values illustrated with asterisks are ** < 0.01,*** <0.001, **** <0.0001. Error bars are mean ± SEM.

Mechano-responses to substrate rigidity by other cell types during adhesion and migration involve microtubule acetylation, which promotes the release of GEF-H1 from microtubules into the cytoplasm to increase Rho activity and cell contractility (Seetharaman *et al*., 2022). We therefore evaluated levels and localization the Rho-GTPase GEF-H1, which is released from acetylated microtubules upon BCR activation and interacts with the exocyst complex to promote lysosome tethering at the immune synapse (Sáez *et al*., 2019). By using the experimental setup described above, we observed that B cells interacting with stiffer substrates, displayed higher levels of GEF-H1 at the IS plane compared to cells activated on soft substrates (Fig EV2 E-F). This result is consistent with B cells progressively enhancing the acetylation of tubulin at the IS interface upon stimulation with stiffer substrates. Collectively, these findings strongly support the notion that B cells respond to physical cues by enhancing tubulin acetylation, which is coupled to the accumulation of GEF-H1 at the synaptic membrane, previously shown to promote lysosome tethering at this interface. Therefore, we next evaluated whether this mechanosensory response directly impacts lysosome trafficking at the immune synapse of B cells.

### Substrate stiffness regulates lysosome positioning and dynamics at the IS

To assess whether physical cues sensed by B cells have an impact on lysosome positioning, we seeded B cells on soft or stiff substrates containing BCR^+^ ligands, at increasing time points. Cells were then stained for the lysosomal marker LAMP1 together with an actin cytoskeleton marker and imaged using confocal microscopy (Fig. 3 A). The distribution of LAMP1^+^ vesicles within the intracellular space, defined by cortical actin cytoskeleton was analyzed as follows: the total cell area was divided into 3 equally concentric sections (Fig. EV2 G) and raw intensity fluorescence values obtained were normalized with respect to the spreading area of each cell. As shown in Fig. 3B, cells stimulated on stiffer surfaces exhibited increased accumulation of lysosomes at the IS, after 15 minutes of activation compared to softer substrates. The amount of lysosomes at the center of the IS was comparable in both conditions after 30 minutes of activation. However, the number of lysosomes at the cell periphery was lower in B cells interacting with stiffer substrates, suggesting that the lysosomes are retained more efficiently at the center of the synaptic plane in B cells activated by antigens associated to surfaces with higher stiffness (Fig. 3B).

**Fig. 3.**
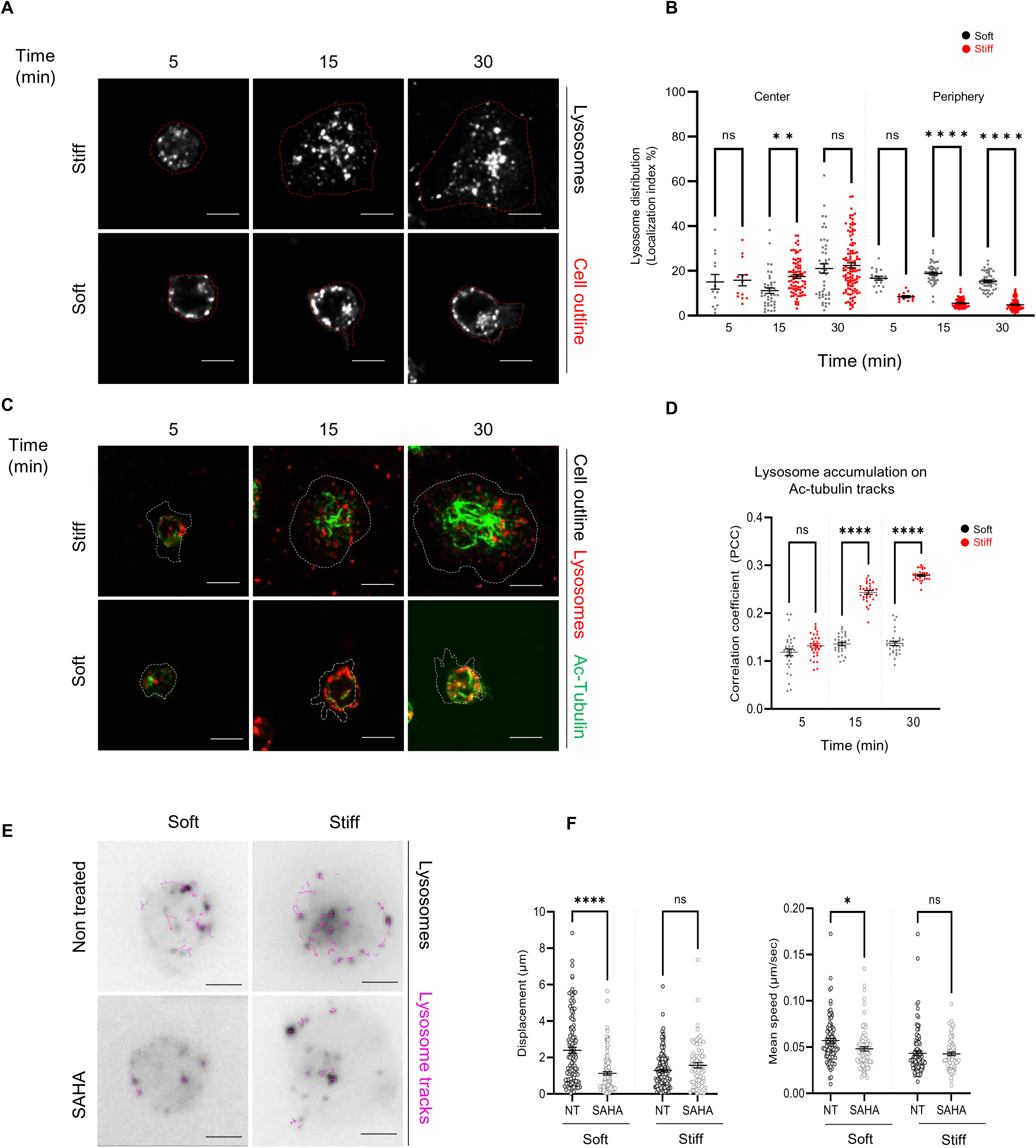
Lysosome localization and dynamics is dependent on tubulin acetylation as a mechano-response of B cell activation (A) Representative confocal images of B cells seeded over stiff or soft substrates at different activation times. Cells were fixed and stained for LAMP1 and actin. The cell border (red outline) was defined by actin phalloidin staining. (B) Quantification of lysosome (LAMP1) accumulation either in the IS center or periphery for cells in (A). The localization index reflects the MFI in a defined area with respect to the total staining in the cell. (C) Representative images of cells seeded in stiff or soft substrates at different time points. The cell outline was determined with actin staining (white dashed lines). The synaptic plane is shown with stainings for lysosomes (red) and acetylated microtubules in green. (D) Quantification of lysosome accumulation on acetylated microtubule tracks from cells shown in (C). Pearson’s correlation coefficient (PCC) is shown for the overlap of both stainings. (E) Representative timelapse images showing the tracking lysosomes stained with lysotracker (black) on either stiff or soft conditions control (Non treated – NT) cells and SAHA-treated cells. Lysosome tracks were followed for 5 minutes after seeding the cells for 15 minutes. Tracks in magenta represent the 15-20 minutes time frames of interaction with the different substrates. (F) Displacement and mean speed of lysosome tracks for experiments detailed in (E). Data shown considers n ≥ 30 cells pooled from N = 3 biological replicates. All scale bars are 5 μm. P values illustrated with asterisks are * < 0.05, ** < 0.01, **** <0.0001. Error bars are mean ± SEM.

Additionally, we quantified the number of LAMP1^+^ particles at the z-plane juxtaposed to the antigen-coated surface (Fig. EV3 A.) in B cells interacting with soft or stiff substrates. As expected, we found more lysosomes at the IS plane of B cells activated onto stiffer substrates. Interestingly, B cells on stiffer gels and increasing activation times displayed smaller lysosomes compared to cells activated on softer substrates (Fig. EV3 B). This could result from changes in lysosome fusion or fission dynamics mediated by mechanical cues that remain to be explored. We next evaluated how substrate stiffness sensed by B cells regulates lysosome dynamics. To this end, B cells labelled with lysotracker were seeded on soft or stiff substrates coupled to BCR^+^ ligands and live imaging was performed for 5-20 minutes, an adequate time frame to evaluate lysosome dynamics during the IS formation (Sáez *et al*., 2019). Consistent with results obtained in fixed cells, lysosomes preferentially accumulated to the cell center upon activation under stiff conditions, whereas in softer substrates, lysosomes were evenly dispersed (Fig. EV3 C). When comparing the mean speed and displacement of lysosomes at the IS, we found that in B cells seeded onto stiffer substrates, both values were lower compared to softer ones (Fig. EV3 D), confirming that lysosome motility and positioning are tuned by extracellular physical cues during B cell activation.

To investigate the functional impact of MT acetylation on lysosome dynamics, we first evaluated the co-localization of Ac-tubulin and LAMP1 in B cells activated on soft and stiff substrates. Our results show that in B cells seeded on stiff substrates, lysosomes progressively increased their co-localization with Ac-tubulin (Fig.3 C-D), whereas this parameter remained constant in B cells activated on soft substrates. These results suggest that there is a functional link between microtubule acetylation during B cell mechano-responses and the positioning of lysosomes at the IS during B cell activation.

To precisely determine the effect of tubulin acetylation on lysosome dynamics in B cells, we evaluated the effect of SAHA, which enhances tubulin acetylation by inhibiting the deacetylase enzyme HDAC6 (Sáez *et al*., 2019). Treatment with 1 μM SAHA for 30 min effectively led to enhanced MT acetylation in B cells (Fig. EV3 E-F). Next, lysotracker-labeled cells were activated on stiff and soft surfaces for 15 min and imaged during 5 min at 3 seconds per frame. Lysosome trajectories from each cell under different conditions were acquired and analyzed. Our results show that lysosomes from SAHA-treated B cells activated on soft substrates, exhibited lower mean speed and were more confined compared to lysosomes from non-treated cells, displaying parameters similar to lysosomes from B cells activated on stiff substrates (Fig. 3 E-F). Interestingly, treatment with SAHA had no significant effect on lysosome dynamics in B cells activated on stiff substrates, indicating that the level of tubulin acetylation or the threshold of tubulin acetylation to control lysosome mobility and positioning achieved by B cells activated on stiff substrates cannot be further increased by inhibiting HDAC6. Overall, this result suggests that tubulin acetylation directly impacts lysosome dynamics in B cells as a response to extracellular stiffness.

### ATAT1 translocates to the Cytoplasm of B cells activated on stiff surfaces

The impact of tubulin acetylation on lysosome dynamics prompted us to investigate the regulatory mechanisms controlling this process. Tubulin acetylation is catalyzed by the acetylase enzyme ATAT1, which shuttles between the nucleus and cytoplasm (Deb Roy *et al*., 2022). Given that tubulin acetylation is enhanced in response to substrate stiffness during B cell activation, we explored whether this was coupled to changes in the localization of ATAT1. To address this, we seeded B cells in soft and stiff substrates and evaluated the localization of ATAT1 by confocal microscopy after different time points of activation. With increasing times of activation and stiffness, ATAT1 shifts from the nucleus to the cytoplasm, which was not observed in B cells activated on soft substrates (Fig. 4 A,B), supporting our hypothesis that BCR-mediated mechanosensing enhances tubulin acetylation by promoting the cytoplasmic localization of ATAT1. To confirm that B cells elicit mechanosensory responses, we show that YES-associated-protein (YAP), a canonical marker of mechanosensing, is translocated to the nucleus in B cells activated on substrates with higher stiffness (Fig. 4 C,D) similarly to other cell types (Panciera *et al*., 2017).

**Fig. 4.**
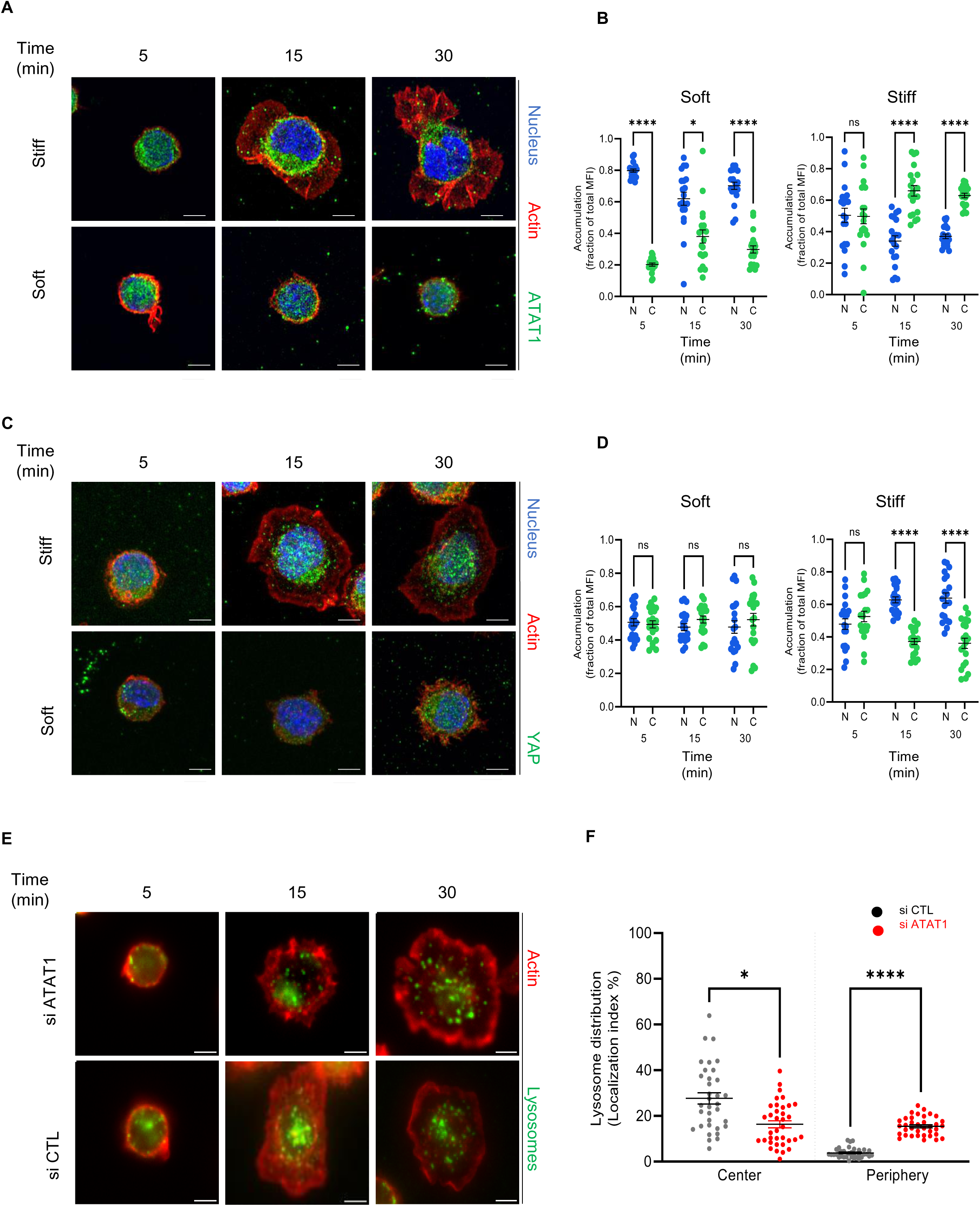
ATAT1 Localizes to the cytoplasm of B cells activated on stiff surfaces to promote microtubule acetylation and regulate lysosome dynamics (A) and (C) Representative images of fixed B cells seeded over stiff or soft substrates at different activation times. ATAT1 is shown in green, the nucleus in blue (Hoechst), and the actin cytoskeleton in red, stained with Phalloidin. (B) Quantification of nuclear or cytoplasmic accumulation of ATAT1 from cells in (A). The distribution index (y-axis) indicates the fraction of ATAT1 fluorescence in the nucleus (N) or the cytoplasm (C, excluding the nucleus) with respect to the total area of the cell. (C) YAP is shown in green, the nucleus in blue (Hoechst), and the actin cytoskeleton in red, stained with Phalloidin. (D) Quantification of YAP nuclear or cytoplasmic accumulation from cells in (C) using the distribution index. (E) Images of B cells seeded over stiff substrates at different activation times in non-silenced (si CTL) or silenced for ATAT1 (si ATAT1). Lysosomes (LAMP1) are shown in green and the actin cytoskeleton in red, stained with Phalloidin. (F) Quantification of LAMP1 accumulation at the IS center or periphery from cell in (E). Shown data considers n ≥ 30 cells pooled from N = 3 biological replicates. All scale bars are 5 μm. P values illustrated with asterisks are: * < 0.05, **** <0.0001. Error bars are mean ± SEM.

### ATAT1 regulates lysosome dynamics

To formally show that ATAT1-dependent microtubule acetylation regulates lysosome dynamics in B cells, we silenced ATAT1 expression using siRNA and evaluated its effect over tubulin acetylation and lysosome distribution. B cells activated on soft substrates display low levels of tubulin acetylation, therefore, to confirm the effect of ATAT1 silencing, we quantified microtubule acetylation in B cells activated on stiff substrates. To this end, B cells were seeded over stiff plastic plates coupled to BCR^+^ or BCR^-^ ligands for 30 minutes. As expected, levels of acetylated tubulin were dramatically reduced upon silencing of ATAT1 compared to control conditions (Fig. EV4 A-B), confirming the role of ATAT1 in this process. Interestingly, cells activated over stiff substrates exhibited greater tubulin acetylation when in contact with BCR^+^ ligands than in control (BCR^-^) conditions (Fig. EV4 A-B). These findings provide additional evidence for the mechanosensing role of the BCR and its link to tubulin acetylation upon its stimulation.

We next analyzed the effect of silencing ATAT1 in lysosome positioning in B cells forming an immune synapse. To this end, B cells were labeled for LAMP1 in ATAT1-silenced and control cells (Fig. 4E, Fig. EV4 C-D) and their distribution at the IS upon stimulation in stiff or soft substrates was analyzed. As anticipated, ATAT1-silenced B cells activated onto stiff substrates, were unable to accumulate lysosomes at the center of the IS after 30 minutes (Fig. 4F). Silencing ATAT1 also decreased pFAK levels (Fig. EV4 E-F) at later time points of activation in B cells seeded onto stiff substrates, suggesting an interplay between the formation of nascent focal adhesions and tubulin acetylation, which could involve the release of actin-factors such as GEF-H1 from the lumen of microtubules when they are acetylated.

Overall, these results reveal that impairing tubulin acetylation by silencing ATAT1 results in defective lysosome positioning at the IS. Hence, ATAT1-mediated tubulin acetylation emerges as a mechanism for controlling the correct positioning and dynamics of lysosomes at the IS in response to external physical cues.

### The efficiency of antigen presentation by B cells depends on substrate stiffness and ATAT1

A fundamental step in B cell activation is the capacity of these cells to present external antigens as peptides loaded on MHC class II molecules to T cells. This enables B-T cooperation and provides survival signals for B cells to sustain their maturation and differentiation into plasma cells (Akkaya, Kwak and Pierce, 2020). To directly evaluate whether physical cues regulate the capacity of B cell to extract and present antigens, B cells were seeded over substrates of different stiffness containing BCR^+^ ligands together with the LACK antigen from *Leishmania major*. The ability of the cells to present MHC-II–LACK processed peptide complexes derived from the substrates to a specific T cell hybridoma that recognizes processed antigen was measured by monitoring IL-2 secretion by activated T lymphocytes. Importantly, we included a positive control for extreme stiffness (glass) coupled to BCR^+^ or BCR^-^ ligands.

As shown in Fig.5 A, B cells seeded on immobilized antigens associated to stiffer substrates, triggered higher IL-2 production from T cells in comparison with B cells activated on soft substrates. This suggests a positive correlation between substrate stiffness, the presence of an activating BCR ligand, and the capacity of B cells to extract, process, and present antigens to T cells. Increasing levels of stiffness had no effect on the presentation of the LACK ^156-173^ peptide, showing that varying substrate rigidity does not significantly influence T cell responses or B-T cell interactions (Fig. 5 B).

**Fig. 5.**
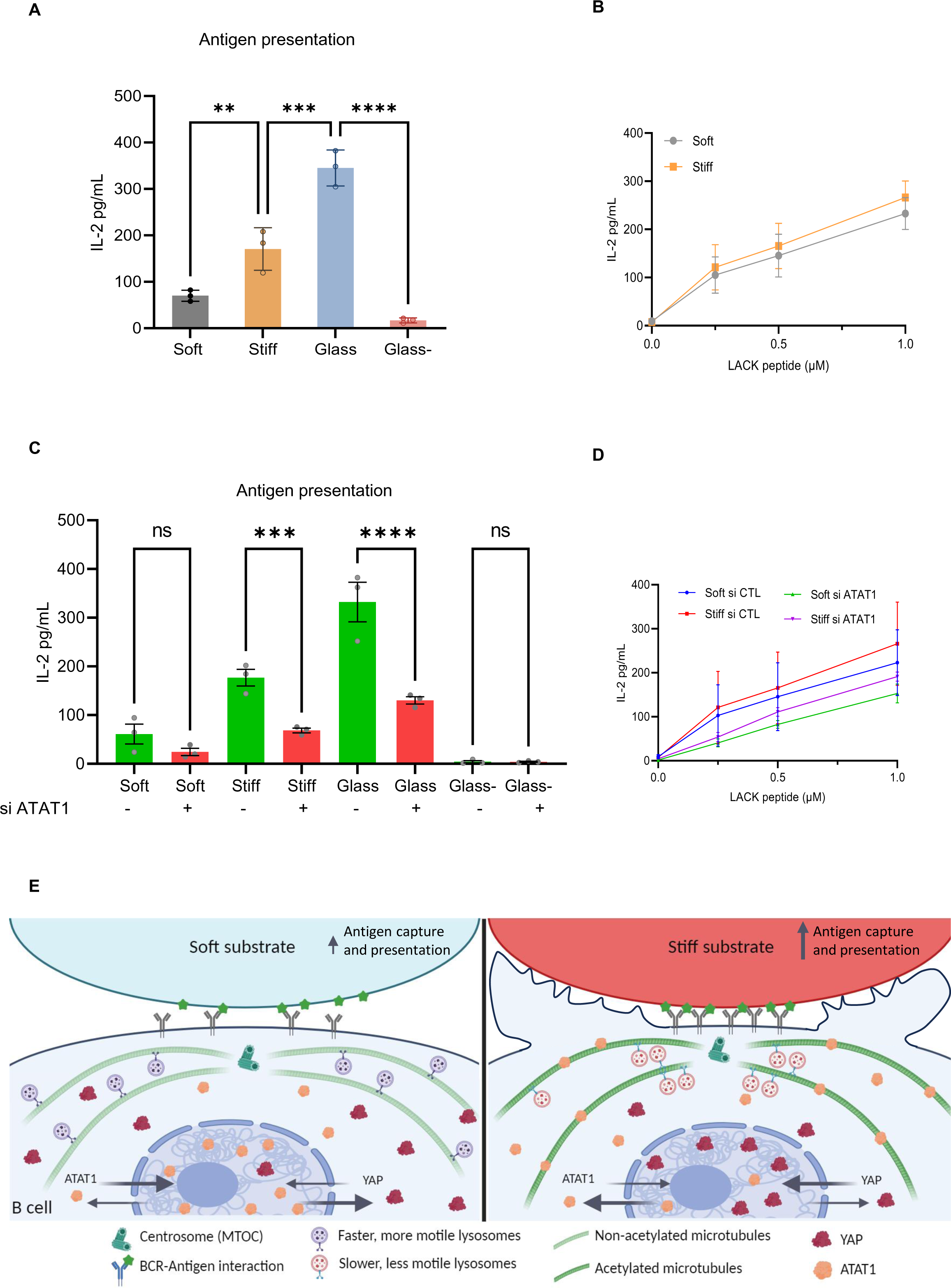
Antigen extraction and presentation by B cells is dependent on the substrate stiffness and ATAT1 Antigen presentation assays (A) of B cells activated on substates with different stiffness and (A) (C) with control or ATAT1-silenced B cells. (B) and (D) Lack peptide control for the cells used. Mean amounts of IL-2 are shown for a representative of three independent experiments performed in triplicate. (E) Proposed model for B cell activation coupled to physical cues: In response to antigens associated to stiff substrates, B cells undergo enhanced spreading responses, increase the formation of actin foci and localize YAP to the nucleus, whereas the acetylase ATAT1 is translocated from the nucleus to the cytoplasm where it catalyzes tubulin acetylation. This enables the central accumulation of lysosomes that preferentially bind to acetylated microtubules at the immune synapse of B cells. These lysosomes exhibit slower speed and displacement than lysosomes from B cells activated on soft substrates. Consequently, B cells interacting with antigens coupled to stiff substrates increase their capacity to capture and present antigens to T cells which is impaired upon ATAT1 downregulation. P values illustrated with asterisks are ** < 0.01,*** <0.001, **** <0.0001. Error bars are mean ± SEM.

Having shown that physical cues from the environment regulate the capacity of B cells to extract and present immobilized antigens, we next evaluated whether this relied on ATAT1-dependent microtubule acetylation. For this purpose, B cells were silenced for ATAT1 and their capacity to extract and present antigens was assessed using the aforementioned experimental setup. As shown in Fig. 4 C, the capacity to present *Lack* antigen associated to stiff substrates was lower in ATAT1-silenced B cells compared to controls cells. However, no significant differences between control and ATAT1-silenced cells were observed when activated on soft substrates. This result is consistent with our observations showing that B cells activated on soft substrates display low levels of translocation of ATAT1 to the cytoplasm and therefore silencing of this enzyme should not have a major effect on antigen presentation. Additionally, the capacity of ATAT1-silenced B cells to present processed LACK^156-173^ peptide was slightly decreased compared to control cells (Fig.5 D), suggesting that B-T cell interactions could be affected. Collectively, these results suggest that silencing ATAT1 renders B cells insensitive to substrate rigidity during activation, compromising their capacity to enhance antigen extraction and presentation.

In summary (Fig. 5 E), our results highlight how B cells couple mechanosensing pathways to control lysosome dynamics and tune their capacity to extract and present immobilized antigens. We show that B cells respond to substrate stiffness translocating ATAT1 from the nucleus to the cytoplasm, thereby increasing tubulin acetylation, which in turn promotes stable lysosome recruitment at the IS. Notably, upon interaction of B cells with stiffer substrates, their lysosomes exhibit a more concentrated pool at the center of the immunological synapse, displaying slower speed and less displacement compared to lysosomes from B cells activated on softer substrates. These changes in lysosome dynamics correlate with the levels of tubulin acetylation, upregulated by B cells in response to substrate stiffness, where lysosomes preferentially associate to acetylated tubulin tracks in stiffer conditions. Overall, these findings underscore the existence of a cellular pathway that connects antigen-extraction mechanisms with mechanical cues originating from APCs, which are regulated by BCR-mediated mechanosensing.

Fig EV1

(A) illustration of basic workflow with PAA gels. First, soft or stiff gels are coupled to BCR-activating ligands. Then cells are seeded for different time points, fixed and used for live imaging, or lysed for protein extraction. (B) Atomic force measurements for stiffness of the gels used in this work. Soft values are considered to be 0.3 kPa and stiff, 13 kPa. (C) representative confocal image for the Z cross-section of a PAA gel coupled to a fluorescently labeled BCR^+^ ligand. Scale bar = 2μm. Bottom: analysis for antigen distribution (BCR^+^ ligand) based on MFI values acquired in confocal microscopy. (D) Representative images obtained in confocal microscopy depicting the actin cytoskeleton spreading labeled with phalloidin (red) when B cells are seeded over gels coupled to a BCR^-^ ligand (non-activating). (E) Quantification of the spreading area from experiments depicted in (D). (F) Immunoblots for pFAK and GAPDH in cells seeded at different time points and stiffnesses. (G) Quantifications for (F) based in the relative abundance of pFAK with respect to GAPDH. Shown data considers n ≥ 30 cells pooled from N = 3 biological replicates. Scale bars are 5 μm. P values illustrated with asterisks are: * < 0.05, ** < 0.01, *** <0.001, **** <0.0001

Fig. EV2

(A) Immunoblot for pAKT, pERK and GAPDH in cells seeded at different time points and stiffnesses. (B) Quantifications for (A) based in the relative abundance of pAKT and pERK with respect to GAPDH. (C) Immunoblotting for Ac-Tub and GAPDH in cells seeded at different time points and stiffnesses. (D) Quantifications for (C) based in the relative abundance of Ac-Tub with respect to GAPDH. (E) Representative images of fixed cells interacting with soft or stiff substrates at different time points. GEF-H1 is shown in a gradient of magenta according to fluorescence intensity. (F) Quantification of GEFH1 intensity at the IS for the experiments represented in (E). (G) Illustration for the localization index used to determine the differential accumulation of lysosomes in each cell. Shown data considers n ≥ 30 cells pooled from N = 3 biological replicates. Scale bars are 5 μm. P values illustrated with asterisks are: * < 0.05, ** < 0.01, *** <0.001.

Fig. EV3

(A) Quantification for LAMP1^+^ particles at the IS showing the number of particles detected at the IS in the different experimental conditions. In (B), every graph depicts summary measurements for individual cells (individual circles) considering the mean area values for segmented LAMP1 particles (x-axis) and the number of segmented particles found at the IS on that cell. (C) Representative images for lysosome tracking in B cells stained with lysotracker (magenta) and activated on either stiff or soft conditions. (D) Displacement and mean speed of lysosome tracks for experiments in (C). (E) Immunoblot to evaluate the effect of the drug SAHA on microtubule acetylation with its correspondent quantification in the fold of change for tubulin acetylation compared to non-treated cells (CTL). (F) Quantification of the effect of SAHA on Ac-tubulin fold of change with respect to total tubulin. Shown data considers n ≥ 30 cells pooled from N = 3 biological replicates. Scale bars are 5 μm. P values illustrated with asterisks are: * < 0.05, **** <0.0001

Fig. EV4

(A) Immunoblot for Acetylated tubulin and GAPDH in cells seeded on stiff plastic plates for 30 minutes. Cells were treated with control or ATAT1 siRNAs. (B) Quantifications for (A) based on the relative abundance of Acetylated tubulin with respect to GAPDH. (C) Representative images of Ac-Tub in control or ATAT1-silenced B cells activated on stiff substrates. (D) Quantification of mean fluorescence intensity of Ac-Tub at the IS as shown in (C). (E) Immunoblot for pFAK and GAPDH in B cells seeded in stiff plastic plates at different time points, for control or ATAT1-silenced cells. (F) Quantifications for (E) based on the relative abundance of pFAK with respect to GAPDH for the treatments at different time points. . Shown data considers n ≥ 30 cells pooled from N = 3 biological replicates. All scale bars are 5 μm. P values illustrated with asterisks are ** < 0.01, **** <0.0001. Error bars are mean ± SEM.

